# Viruses in deep-sea cold seep sediments harbor diverse survival mechanisms and remain genetically conserved within species

**DOI:** 10.1101/2023.03.12.532262

**Authors:** Yongyi Peng, Zijian Lu, Donald Pan, Ling-Dong Shi, Zhao Zhao, Qing Liu, Chuwen Zhang, Kuntong Jia, Jiwei Li, Casey R.J. Hubert, Xiyang Dong

## Abstract

Deep sea cold seep sediments have been discovered to harbor novel, abundant, and diverse bacterial and archaeal viruses. However, little is known about viral genetic features and evolutionary patterns in these environments. Here, we examined the evolutionary ecology of viruses across active and extinct seep stages in the area of Haima cold seeps in the South China Sea. Diverse antiviral defense systems in 487 microbial genomes spanning 43 families inform the dynamics of host-virus interactions. Accordingly, a total of 338 viral operational taxonomic units are identified and linked to 36 bacterial and archaeal phyla. Cold seep viruses are predicted to harbor diverse adaptive strategies to persist in this environment, including counter-defense systems, reverse transcriptases, auxiliary metabolic genes, and alternative genetic code assignments. Extremely low nucleotide diversity is observed in cold seep viral populations, being influenced by factors including microbial host, sediment depth and cold seep stage. Most cold seep viral genes are under strong purifying selection with trajectories that differ depending on whether cold seeps are active or extinct. This work sheds light on the understanding of environmental adaptation mechanisms and evolutionary patterns of viruses in the sub-seafloor biosphere.

## Introduction

Cold seeps are deep sea environments where hydrocarbon fluids and gas seepage occur at the continental margins worldwide. The continuous seepage of gaseous and liquid hydrocarbons boosts local biodiversity and microbial activity, featuring prevalent archaeal anaerobic methanotrophs (ANME) and sulfate-reducing bacteria (SRB)^1, 2^. Compared to the rich knowledge of cold seep bacterial and archaeal communities, viruses remain largely underexplored in spite of their significant roles in impacting microbes and corresponding biogeochemical cycles^3, 4^. Virus studies using enumeration or cultivation have shown that cold seep sediments are hotspots of viral production with high virus-prokaryote ratios^5, 6^. A recent survey of metagenomes from seven cold seeps found that these sediments harbor diverse and novel viruses, hinting at their potential impact on hydrocarbon biodegradation and other local metabolisms catalyzed by cold seep microbiomes^7^. However, cold seep viral diversity and distribution patterns, virus-microbe interactions, adaptive mechanisms to environmental factors, and viral genetic diversity are still relatively unexplored.

Viruses have a genetic toolbox of diverse mechanisms to adapt to the environment and co-evolve with hosts. As foreign mobile genetic elements, viruses face a wide repertoire of antiviral defense systems, including restriction-modification (RM) and CRISPR-Cas^8^. In line with antagonistic co-evolution of viruses and their hosts^9, 10^, viruses have developed efficient and robust counter-defense systems, such as anti-restriction, anti-CRISPR and other counter-defense proteins^11, 12^. Diversity-generating retroelements (DGRs) containing reverse transcriptase (RT) are another important diversification mechanism for driving sustained amino acid-level diversification of their target domains^13, 14^. Viruses also encode DGRs to produce many mutations in specific regions of host target genes through error-prone reverse transcription^15–17^. To replicate more efficiently, viruses can alter their hosts’ metabolic potential through the expression of auxiliary metabolic genes (AMGs) to modulate host cell metabolism during infection^18^. In addition to these gene inventories, viruses can use alternative genetic codes different from those of their host, potentially increasing viral adaptability (e.g., in regulating translation of lytic genes)^19, 20^. Whether or not cold seep viruses incorporate these strategies into their repertoire of mechanisms for mediating host-virus interactions and environmental adaptation in these harsh deep sea subseafloor environments requires further investigation.

Intra-population genetic variations (i.e., microdiversity) can also improve virus adaptation to their environment by driving phenotypic variation^21, 22^. For example, depth-dependent evolutionary strategies of viruses were observed in the Mediterranean Sea^9^ and grassland soil in northern California^10^. Large viral microdiversity was observed for perhaps the most abundant ocean virus in temperate and tropical waters infecting *Pelagibacter*^23^, whereas viruses were under significantly low evolutionary pressures in stable subzero Arctic brines^24^. The principles governing the viral evolution likely differ depending on environmental conditions, such as host dynamics, physicochemical properties, and population sizes^25–27^. Examining 39 abundant microbial species identified in sediment layers below the sea floor and across six cold seep sites, we previously reported that their evolutionary trajectories were depth-dependent and differed across phylogenetic clades^1^. However, it remains to be answered if cold seep viruses are undergoing similar evolutionary patterns and selection pressures.

To understand adaptive survival mechanisms and genetic microdiversity of cold seep viruses, we extracted viral genomes from 16 sediment core samples in the area of Haima cold seeps in the South China Sea (**Supplementary Figure 1 and Supplementary Table 1**). Cores were collected from two active seeps with dense and living bivalves, as well as from one extinct seep covered with many dead clams^28^. We explored viral diversity patterns at both the community-level (i.e., macrodiversity) and population-level (i.e., microdiversity), and viral functional gene repertoire related to arms race between viruses and their prokaryotic hosts. This study expands the knowledge of ecological and evolutionary patterns of viruses inhabiting in cold seep subsurface ecosystems.

## Results

### Diverse antiviral strategies in cold seep microbial genomes

In total, 16 metagenomic data sets were derived from depth-discrete sediment core samples obtained from two active (*n* = 5 for Active1; *n* = 6 for Active2) and one extinct (*n* = 5) cold seeps (**Supplementary Figure 1 and Supplementary Table 1**), at depths ranging from 0 to 20 cm below the sea floor (cmbsf)^28^. Bacterial and archaeal community structures varied between different depth layers at the three sites (**Supplementary Figure 2 and Supplementary Table 2**). Active seep sediments were dominated by Halobacteriota and Desulfobacterota, whereas Desulfobacterota and Chloroflexota were the major microbial lineages in extinct seep sediments. After assembly, 487 species-level metagenome-assembled genomes (MAGs) were reconstructed at an average nucleotide identity (ANI) threshold of 95% (**Supplementary Figure 3 and Supplementary Table 3**), spanning 53 bacterial and 10 archaeal phyla, with the majority affiliated with Proteobacteria (*n* = 59), Desulfobacterota (*n* = 56), Chloroflexota (*n* = 49), Bacteroidota (*n* = 38), and Thermoplasmatota (*n* = 24).

Bacteria and archaea possess diverse antiviral strategies to defend against infection by their viruses^29–31^. A total of 2,145 antiviral genes were detected in 63% of cold seep microbial genomes, and could be assigned to 43 families of antiviral systems^8, 32^; these include Restriction-Modification (RM) systems that target specific sequences on the invading DNA elements and CRISPR-Cas systems that use RNA-guided nucleases to cleave foreign sequences^33^ (**Figure 1a and Supplementary Table 4**). On average, the cold seep microbial genomes encode two antiviral systems per genome and the number of antiviral systems is positively correlated with the genome size for each MAG (linear regression; *R*^2^ = 0.27, *P* = 4.73×10^-5^; **Figure 1b**), similar to previous observations on the importance of genome size for encoding accessory systems in prokaryotes or ocean microbiome^8, 34^. The number of antiviral systems per genome varies from zero (179 genomes) to 32 in a genome belonging to the phylum Fermentibacterota (classified as JAFGKV01 at the family-level; **Supplementary Table 4**), followed by 30 in a *Gammaproteobacteria* genome and 27 in a *Bacteroidia* genome. The most abundant species in the metagenomic dataset (18% of the microbial community) is the putative anaerobic methanotroph ANME-1 SY_S15_40 that encodes two RM type II and one RM Type IIG systems (**Supplementary Tables 3 and 4**). Based on surveying large datasets of sequenced genomes, RM and CRISPR-Cas systems were reported to be present in ~75% and ~40% of microbial genomes, respectively^29, 35^. Relatively fewer cold seep microbial genomes appear to encode RM (50.8%) and CRISPR-Cas systems (22.7%), yet feature higher frequencies of AbiEii (44%; one antiviral system of Abortive infection^36^) and SoFlC (38%) that can modulate various target protein activities^32^ (**Figure 1c and Supplementary Table 4**). Overall, these data reveal diverse antiviral strategies throughout the Haima cold seep microbiome with specific enrichment in some antiviral systems that govern the dynamics of host-virus interactions.

**Figure 1.**
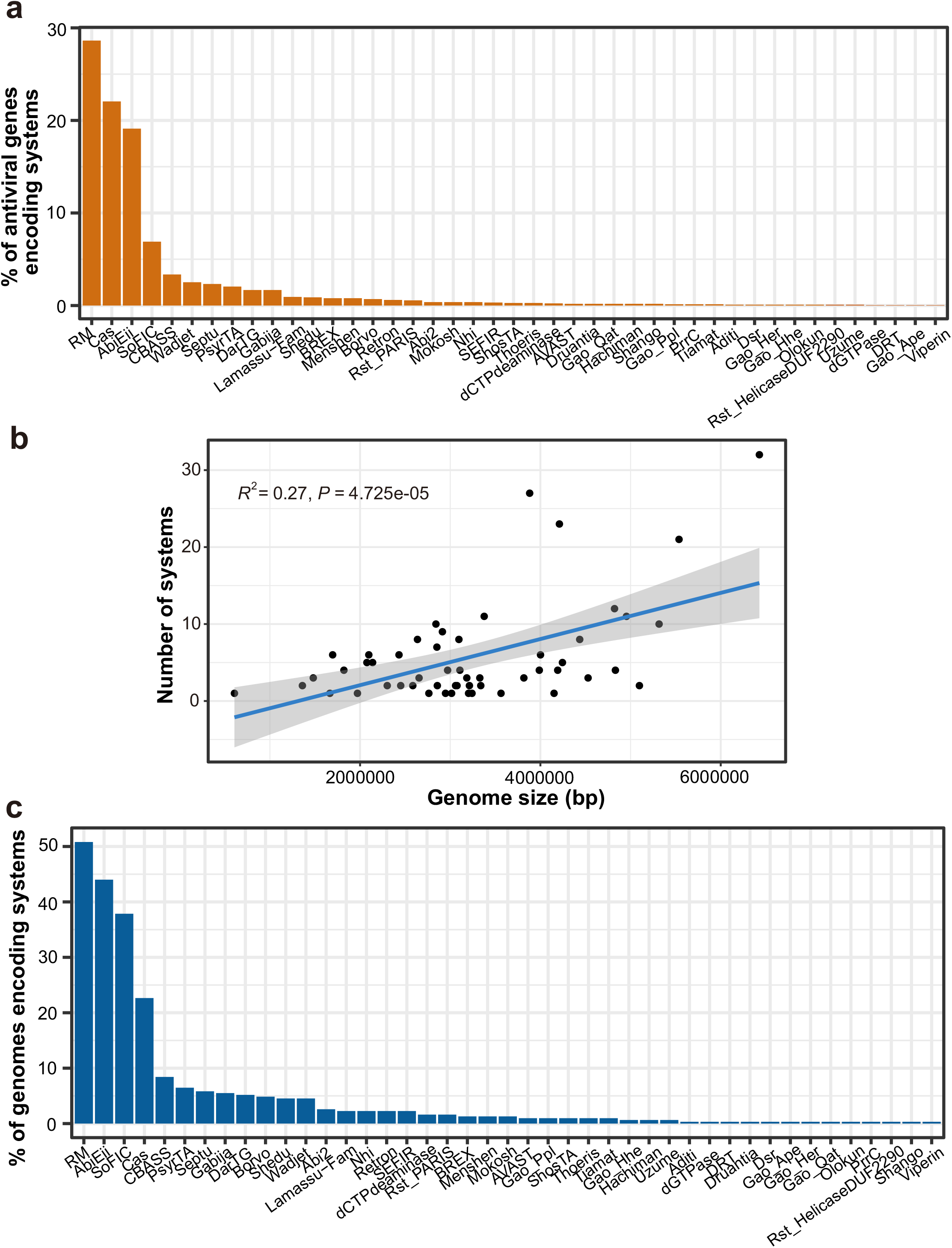
Diversity of antiviral systems found in cold seep bacterial and archaeal genomes. (a) Frequency of genes encoding antiviral systems in all genomes. (b) Relationship between system numbers per prokaryotic genome and genomic size. The correlation analysis was conducted with the completeness-filtered dataset (> 90% completeness) to reduce the potential bias caused by incompleteness. (c) Frequency of antiviral systems detected in all genomes. Detailed statistics for antiviral systems of MAGs are provided in **Supplementary Table 4**.

### Novel viral genomes linked to 36 microbial phyla

Cold seep samples contained highly abundant viruses with densities up to 7.6×10^11^ per gram sediments, with viral abundances being associated with sediment depth (**Supplementary Table 5**). From the 16 metagenomic data set, 488 single-contig viral genomes with ≥50% estimated completeness (based on CheckV^37^) were recovered using multiple virus identification tools (**Figure 2a and Supplementary Figure 4**). Viral genomes were clustered into 338 species-level viral operational taxonomic units (vOTUs)^38^, belonging to 83 viral clusters (VCs; roughly equivalent to an ICTV genus) utilizing whole genome gene-sharing profiles^39^ (**Supplementary Figure 5 and Supplementary Table 6**). Similar to observations in prokaryotic communities^1, 2, 40^, alpha and beta diversity analyses of 338 vOTUs suggest that sampling site, sediment depth in relation to redox conditions^28^, and the geological state of cold seeps (active or extinct) shape the structure of viral communities (**Supplementary Figure 6 and Supplementary Table 5**).

**Figure 2.**
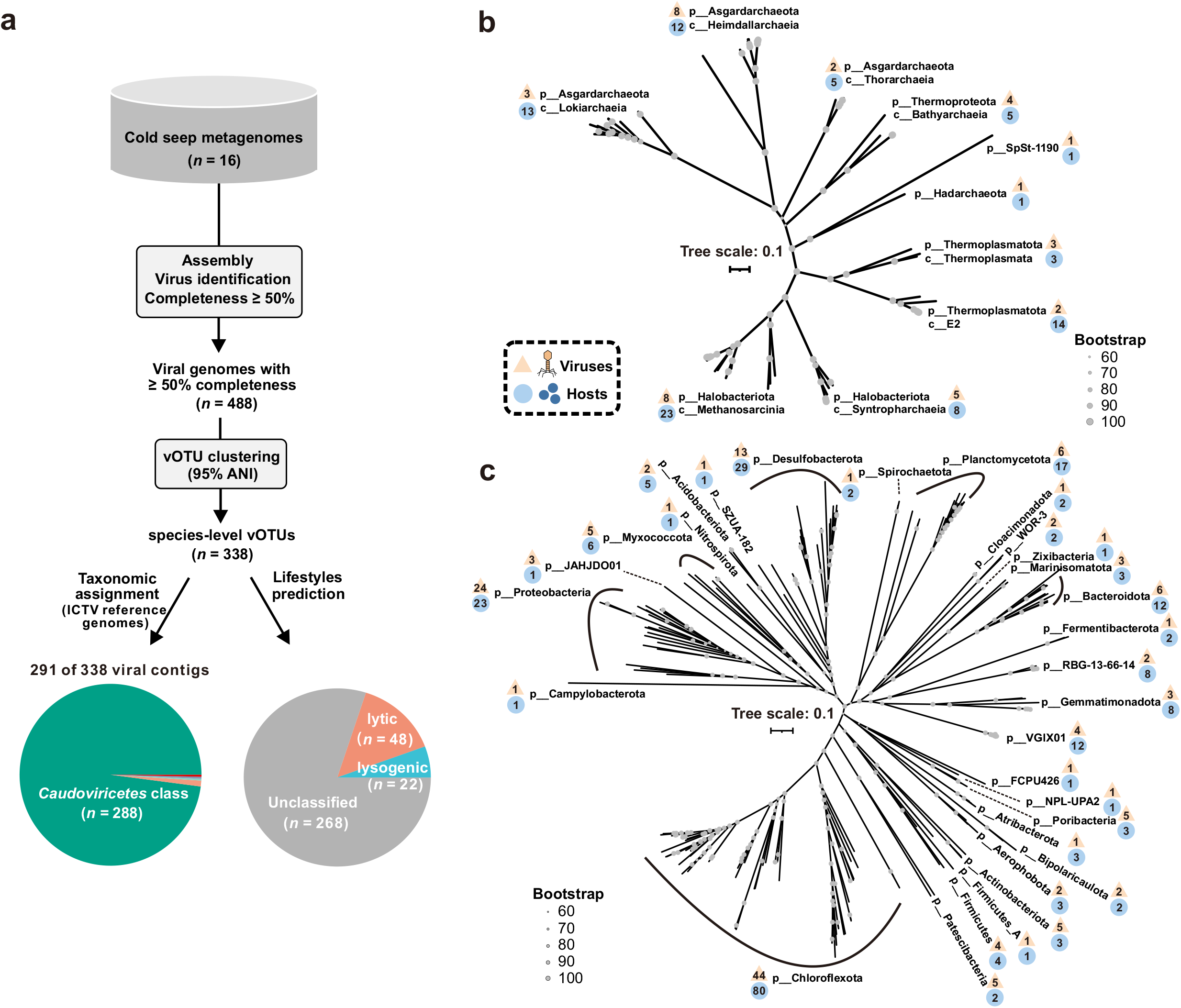
Ecological features of novel cold seep viruses. (a) Workflow for identification, taxonomic assignment and lifestyle prediction of viruses. Phylogenetic trees of predicted hosts for viruses constructed based on marker genes using GTDB-Tk, including archaeal (b) and bacterial (c) phylogenetic trees. Detailed statistics for taxonomy, lifestyles and host-virus linkages of viruses are provided in **Supplementary Tables 6 and 7**.

Among the 338 vOTUs, 291 could be taxonomically assigned revealing that 288 are affiliated with the class *Caudoviricetes* (**Figure 2a and Supplementary Table 6**), which encompasses tailed phages that are the most prevalent viral taxon across ecosystems^41^. Only ten vOTUs could be annotated at the order level, confirming a large knowledge gap in the taxonomy of deep-sea cold seep viruses^7^. With respect to viral lifestyles, 48 and 22 vOTUs were predicted to be lytic and lysogenic, respectively, with others being unclassified (**Figure 2a**). Host predictions of these vOTUs revealed that virus-infected hosts were detected in 36 bacterial and archaeal phyla (**Figures 2b-c and Supplementary Table 7**). From the 475 host-virus linkages, the most common phylum among predicted hosts was Chloroflexota (*n* = 80), followed by Halobacteriota (*n* = 31), Asgardarchaeota (*n* = 30) and Desulfobacterota (*n* = 29). Ten viruses were predicted to infect ANME-1 and ANME-2 groups that perform anaerobic methane oxidation. Viruses infecting *Methanosarcinales* and *Gammaproteobacteria* were highly abundant in the extinct and active cold seep samples, respectively.

### Cold seep viruses harbor diverse strategies for environmental adaptation

To protect against antiviral systems of their microbial hosts, cold seep viruses encode an extensive repertoire of counter-defense systems, including anti-CRISPR (Acr) proteins, methyltransferases and antitoxins (**Figures 3a-c and Supplementary Table 8**). A total of 75 type II DNA methyltransferases without counterpart restriction enzymes were detected in 55 viral genomes, encoding diverse DNA modification enzymes (e.g., adenine- and cytosine-specific methyltransferases and adenine methylase)^34^. The *acr-aca* operon (anti-CRISPR gene *acr* and *acr*-associated gene *aca*)^42^ was identified in 10 viral genomes, which may inhibit the CRISPR-Cas immunity of the host to allow viruses to propagate^43^. Accordingly, one Poribacteria genome SY_Active_Co137 infected by a virus with the *acr*-*aca* operon has 9 *cas* genes (**Supplementary Tables 4 and 8**). Interference modules of the antitoxin genes (e.g., *vapBC, relBE, hicBA*) were found in 63 viruses and belonged to the type II Toxin-antitoxin (TA) system^44^. Additionally, a total of 17 viruses were found to encode two or more types of counter-defense systems. Different classes of reverse transcriptases (RTs) were also found in 22 viruses, including diversity-generating retroelements (DGRs), retrons, UG26 and UG28 (**Figure 3e and Supplementary Figure 7**). Among them, RTs associated with DGRs were detected in five viruses; this mechanism can introduce variations in the target gene and facilitating the evolution of their hosts^17^. Retrons were found in three viruses also possibly involved in defense systems for foreign DNA elements^44, 45^.

**Figure 3.**
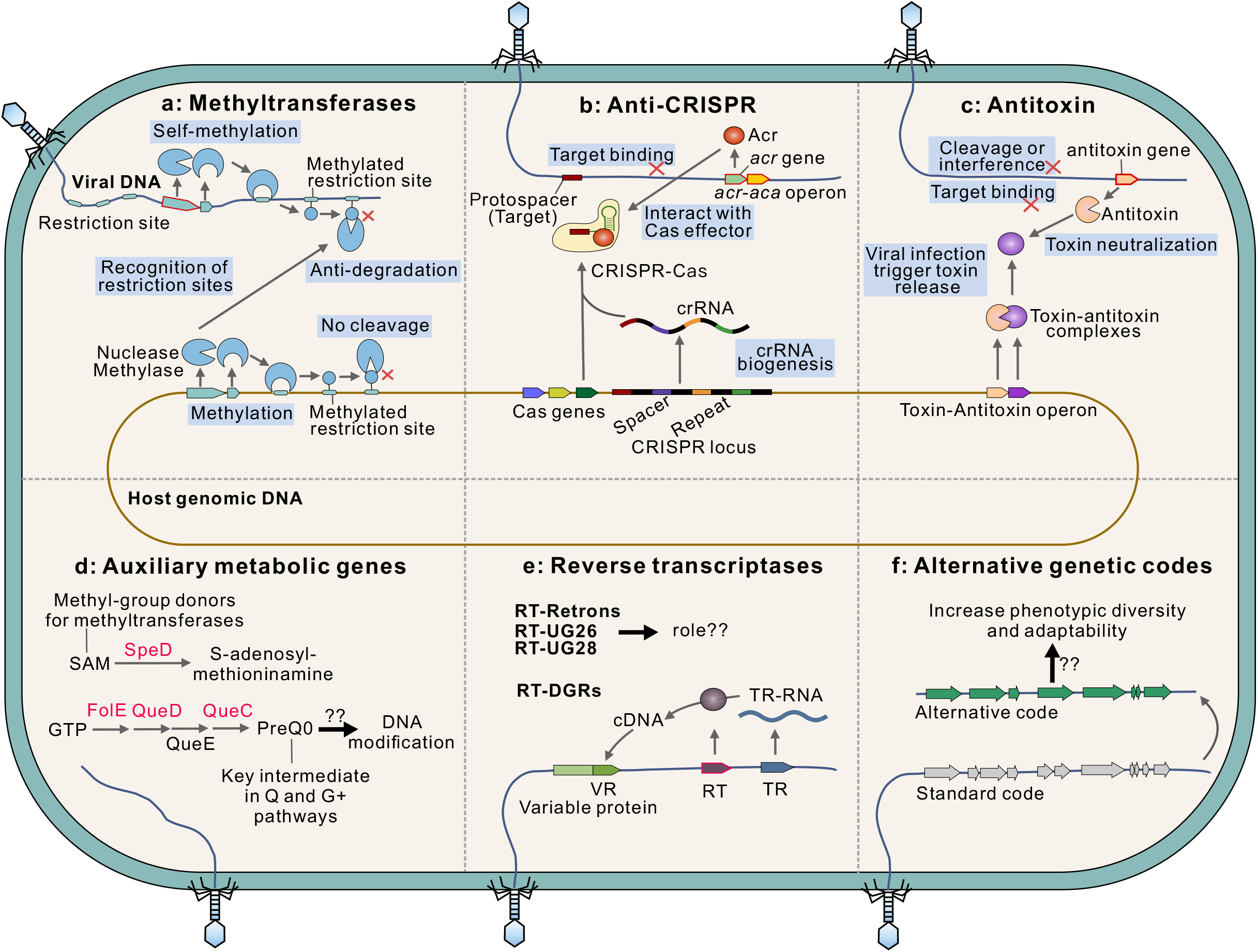
Diverse strategies for environmental adaptation in cold seep viruses. (a) Viruses can encode methylases that modify their DNA to prevent its recognition by host restriction-modification (R-M) systems and cleavage by certain restriction endonucleases. (b) Anti-CRISPR (*acr*) genes encoded on the viruses can inhibit CRISPR-Cas activity when it may have been targeted by the CRISPR-Cas system of the host. (c) Viruses can encode antitoxins that can neutralize host toxin-antitoxin (TA) systems. (d) Potential function for how virus encoded auxiliary metabolic genes (AMGs) contribute to the fitness of the host and/or viruses. SAM (S-adenosylmethionine) is well known as the methyl donor for methyltransferases that modify DNA, RNA, histones and other proteins. Decarboxylation of SAM to S-adenosylmethioninamine may reduce the SAM required for methylation by host enzymes. 7-cyano-7-deazaguanine (preQ0) is synthesized from GTP by four enzymes (FolE, QueD, QueE, QueC) and is the key intermediate in both the Q and G^+^ pathways, which can be further modified for protecting viral DNA from host restriction enzymes. (e) Reverse transcriptases (RTs) including diversity generating retroelements (DGRs), retrons systems, and UG26 and UG28 systems are found in cold seep viruses. Reverse transcriptase (RT) mediates exchange between two repeats: one serves as a donor template (TR) and the other as a recipient of variable sequence information (VR), resulting in massive sequence variation in the receptor-binding protein. Multiple other systems have been identified in viruses, but their roles and mechanisms remain unknown. (f) Alternative genetic codes are seen in the genomes of some cold seep viruses. These have recently been demonstrated to increase adaptability^20, 48^, but their roles in cold seep viruses remain unknown. Related genes identified in cold seep viruses are marked in red (gene name) or with red border (gene arrow). Detailed statistics for diverse strategies for environmental adaptation in viruses are provided in **Supplementary Tables 8-10**.

As an important mechanism in adaptation to the environment, viruses can acquire new functional genes via transduction, i.e., auxiliary metabolic genes (AMGs) that contribute to host and/or viral fitness^4, 46^. Ten AMGs were identified in 7 viral genomes (**Figure 3d, Supplementary Figure 8 and Supplementary Table 9**), related to four different types of functions. Two AMGs encoded GTP cyclohydrolase I (FolE), and six belonging to Que super family (QueC and QueD) may synthesize GTP to 7-Cyano-7-deazaguanine (preQ_0_) for genome modifications and translational efficiency^47^. AMGs encoding S-adenosylmethionine decarboxylase (SpeD) and Dehydrogenase E1 component were also identified, and are involved in biosynthesis of amines or polyamines and the tricarboxylic acid cycle, respectively. Diverse lineages of viruses from different habitats have been seen to be self-beneficially employ alternative genetic codes to reassign one or more codons^20, 48–50^. In the dataset from the Haima cold seeps, 16 viral genomes are predicted to use genetic codes characterized by reassignments of the *ochre* (TAA; *n* = 620 recoding events of genes), *amber* (TAG; *n* = 182) or *opal* (TGA; *n* = 3) stop codons (**Figure 3f, Supplementary Figure 9a and Supplementary Table 10**). These viruses are associated with hosts in multiple phyla (e.g., Desulfobacterota and Acidobacteriota). Genome sizes of these viruses range from 5.2 kb to 179.7 kb, with larger genomes having more recoding events of genes (linear regression; *R^2^*=0.58, *P*=0.0004). Recoded genes were mostly associated with replication, recombination and repair functions, followed by unknown functions (**Supplementary Figure 9b**), suggesting adaptive recoding in controlling viral replication and regulation.

### Cold seep viruses are genetically conserved and under strong purifying selection

Nucleotide diversity (π), single nucleotide polymorphisms (SNPs) and fixation indices (F_ST_) were calculated to track viral microdiversity (**Supplementary Tables 11 and 12**). Nucleotide diversity of cold seep viral populations ranged from zero to 3.06×10^-3^, and were on-average 1.29×10^-4^ (median 3.38×10^-5^) for viruses detected in both active and extinct cold seep sediments (**Figure 4a**). This viral nucleotide diversity is significantly lower than that observed for viral populations in seawater sampled from throughout the world’s oceans (on-average 3.78×10^-4^)^22^ and in soils having various land uses (on-average 6.54×10^-3^)^51^. Low SNP frequencies were also observed in Haima cold seep viral populations (0.33 SNP per 1000 bp on average, median 0.076; **Figure 4b**), e.g., compared to those detected in the SARS-CoV-2 coronavirus, in 25 uncultivated virophage populations in North American freshwater lakes, and in 44 dsDNA viral populations dominating the oceans, based on various approaches for the extraction of viral genomes^52–54^. F_ST_ values between viral populations in relation to different sediment samples ranged from zero to 0.89 and were on-average 0.048, with 80% of pairwise fixation indices being zero (**Figure 4c**). These data reflect that cold seep viral populations are genetically conserved and homogeneous contrary to observations of their microbial hosts^1^, suggesting viruses and microbes might undergo different types of environmental selection.

**Figure 4.**
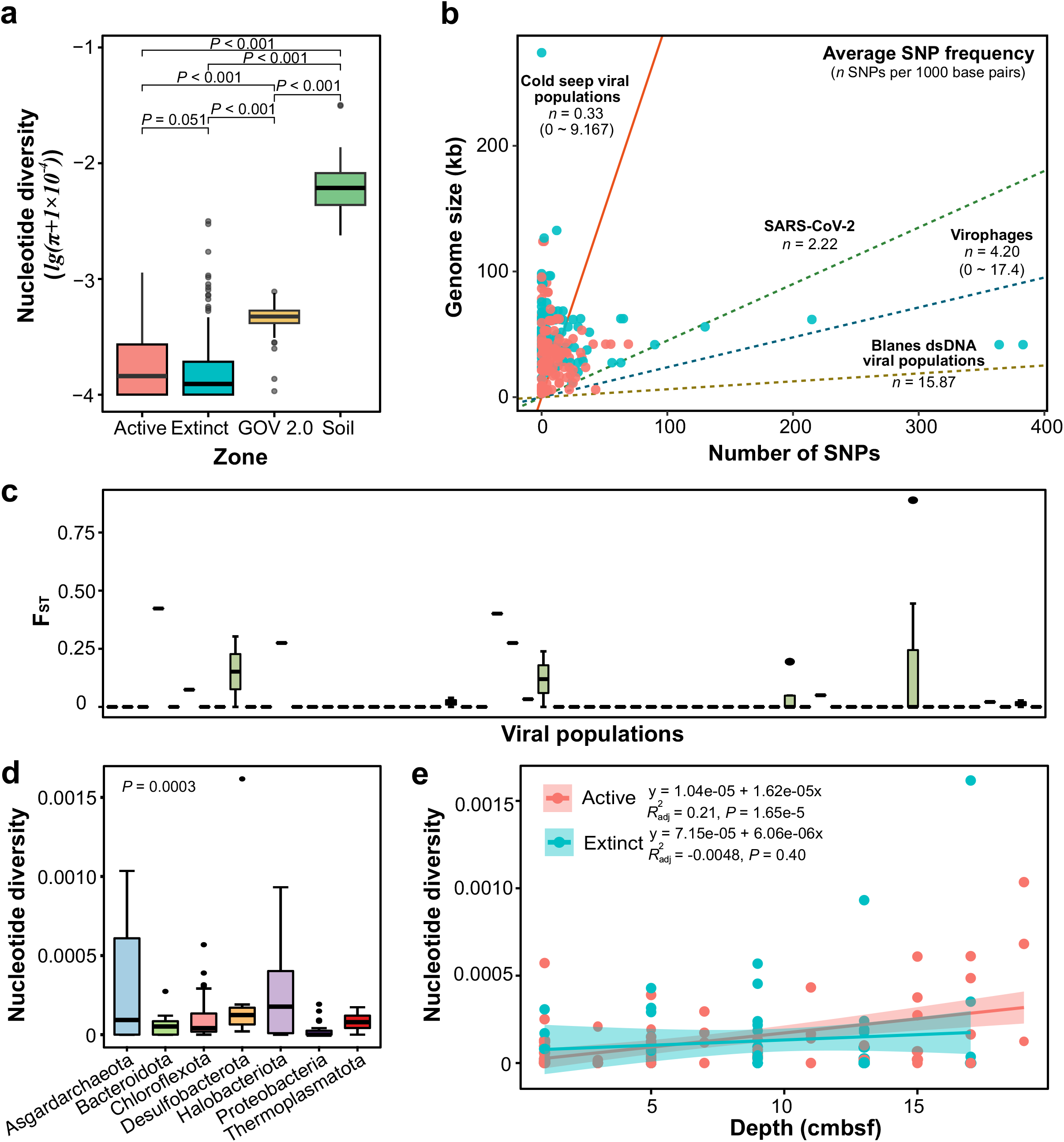
Genome-wide evolutionary metrics of cold seep viral populations. (a) Nucleotide diversity of viruses from active and extinct cold seeps (this study), seawater sampled at a global scale (GOV 2.0) and soil samples from various land-use types. (b) Comparisons of average SNP frequency among cold seeps viruses, SARS-CoV-2, virophages and blanes viruses (dominant dsDNA populations). Linear regressions are indicated for different viral groups. (c) Boxplot showing the F_ST_ values measured as the differences between the same viral populations found in two distinct cold seep samples. (d) Genome-wide nucleotide diversity of viruses infecting different microbial populations. (e) Comparison of nucleotide diversity of viruses infecting dominant populations against sediment depths at the genome level. Linear regressions and *R* values are indicated for viruses in active and extinct cold seeps. Sites color code: blue: extinct cold seep; red: active cold seep. Detailed statistics for evolutionary metrics of cold seep viruses are provided in **Supplementary Tables 11-12**.

Nucleotide diversity of viral populations is significantly different among viruses infecting different microbial hosts (*P*= 0.0003; **Figure 4d and Supplementary Table 11**). Archaeal viruses associated with Halobacteriota have the highest nucleotide diversity. Like evolutionary trajectories of microbial populations in cold seeps^1^ (e.g., Asgardarchaeota, Halobacteriota and Bacteroidota), the nucleotide diversity of associated viruses is also depth-dependent in active cold seeps (**Figure 4e**). On the other hand, no obvious depth-dependent trends were observed for viruses in the extinct cold seep. This is in agreement with the significant difference for nucleotide diversity between the two cold seep stages (**Figure 4a;***P*= 0.051).

At the gene level, 90.6% of pN/pS ratios were less than 0.4, much lower (*P* < 0.0001) than those observed for viral assemblages present in underground saline waters from hypersaline springs^55^ (**Figure 5a, Supplementary Figure 10 and Supplementary Table 13**), indicating that most cold seep viral genes were under strong purifying selection. However, genes under positive selection were also detected in relation to viral DNA replication, recombination, repair and maturation (**Figure 5b**), including *terL*, *polA* and *dnaB* with abnormally high pN/pS values (**Supplementary Table 13**). Significant differences were exhibited for pN/pS ratios between the two cold seep stages (**Figure 5a;** *P* < 0.0001). When grouped according to the functional categories of VOGDB (http://vogdb.org/), nucleotide diversity values were found to be significantly different while no significant differences were observed for pN/pS ratios (**Supplementary Figure 11**). Tajima’s D values ranged from −9.7 to zero and significantly varied (*P* = 1.66×10^-8^) between the two cold seep stages (**Figure 5c**). A total of 90.5% of viral gene Tajima’s D values were found to be zero with no detected SNP. For others, genes under natural selection (Tajima’s D < −2.5; 6.1%) outnumbered those under neutral processes (Tajima’s D = 0; 3.4%). The observation of large number of negative values supports the presence of excess rare alleles and recent expansion of cold seep viral populations^56^.

**Figure 5.**
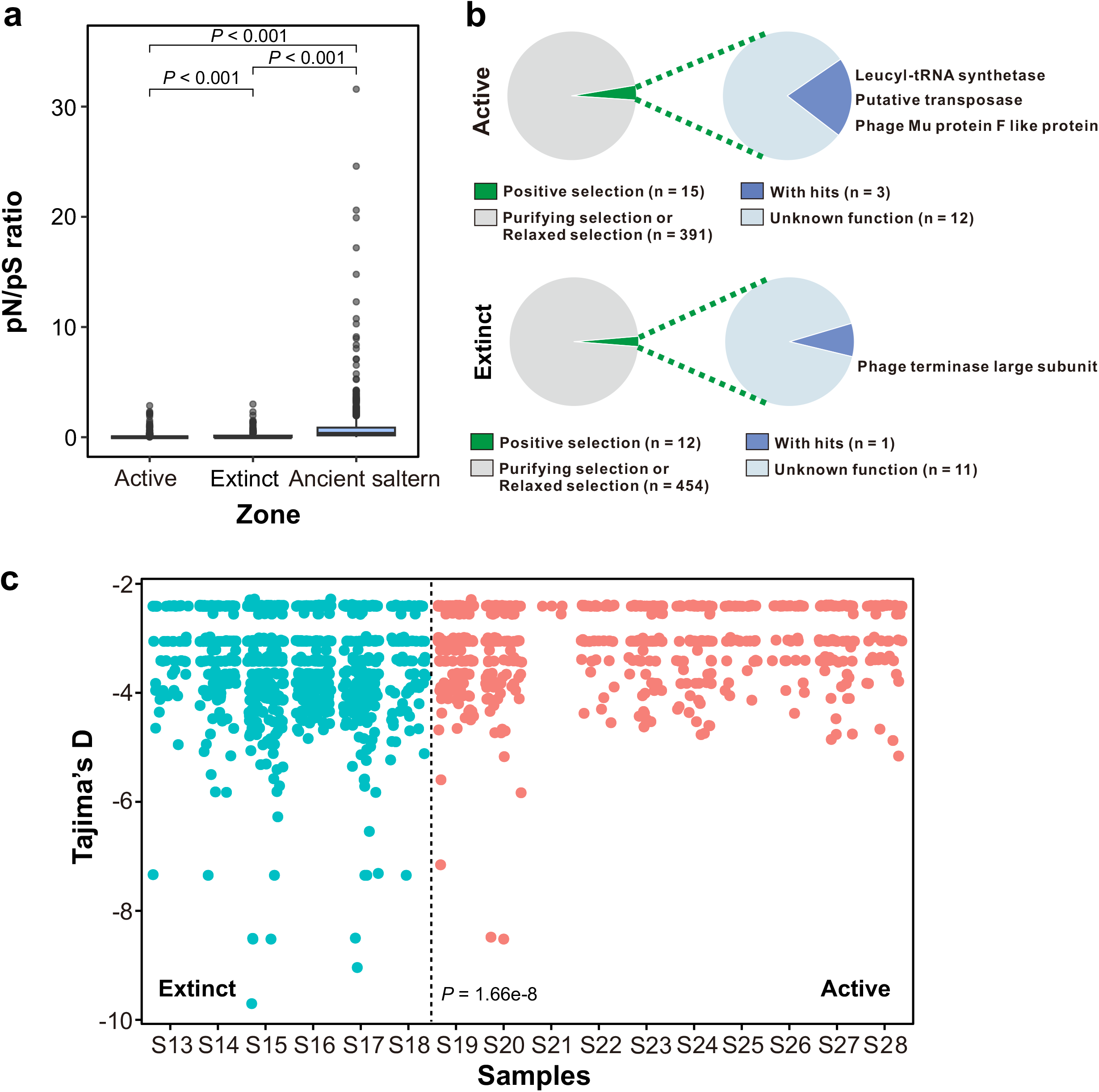
Gene-wide evolutionary metrics of cold seep viral populations. (a) pN/pS ratio of viral genes from cold seeps (this study) and viral genes from an ancient saltern^55^. (b) Viral genes under positive selection in active and extinct cold seeps. Viral genes are further divided into two groups based on pN/pS values, i.e., genes under positive selection (pN/pS ≥ 1) and those under purifying selection or relaxed selection (pN/pS < 1). (c) Tajima’s D of viral genes across 16 sediment samples from extinct (blue) and active (red) cold seeps. Detailed statistics for evolutionary metrics of cold seep viruses are provided in **Supplementary Table 13**.

## Discussion

To date, studies of viral ecology and evolution have paid little attention to how subsurface viruses evolve to adjust to their surrounding environment and interact with their hosts^4, 25, 26, 57^. Besides investigating structural and functional characteristics of viral communities, this study highlights evolutionary adaptation patterns of viruses at different sediment depths in cold seeps that are active and extinct. Novel and abundant deep-sea cold seep viruses were identified and observed to vary between active and extinct cold seeps and different sediment depths. These viruses are associated with major lineages of cold seep archaea and bacteria, including many taxonomic groups with no cultured representatives. Cold seep archaea and bacteria have various antiviral defense systems to prevent infections of diverse and abundant viruses, such as RM, AbiEii, SoFlC and CRISPR-Cas systems. Likewise, their viruses have evolved to harbor a rich repertoire of adaptive strategies to defend against these host systems, including anti-CRISPR (Acr) proteins and methyltransferases. In addition to counter-defense systems to combat microbial hosts, cold seep viruses also contain RTs and AMGs that contribute to viral fitness, as well as alternative genetic code assignments to increase phenotypic diversity. Beyond genetically diverse features of cold seep viruses, their evolutionary trajectories are also surprisingly unique, featuring genetic conservation and homogenous genomes with unexpectedly low microdiversity. Most viral genes generally undergo strong purifying selection, in both the active and extinct cold seep sediments. These findings indicate that multiple factors are likely to determine the evolutionary patterns of cold seep viruses, including microbial hosts, sediment depth and cold seep geology.

Together, these analyses of evolutionary dynamics of viruses will help guide future studies targeting the viral evolution and virus-host systems in extreme environments. However, it should be noted our results are representative only of double-stranded DNA viruses, such that other viral particles are not incorporated in the extraction process and analysis^9^. Nevertheless, studies with more samples from more locations and covering larger spatial gradients via the combination of metagenomes and viromes as well as single-virus genomics^23, 52^ will be necessary to determine if the trends presented here are universal for deep sea subseafloor viral communities.

## Methods

### Sample description, metagenomic sequencing and analysis

Metagenomic sequencing was performed on 16 sediment samples collected from the Haima cold seeps in the northern part of the South China Sea (**Supplementary Figure 1**). Samples were taken from two active seep sites and one extinct seep site via the R/V Tansuo Yihao using the piloted submersible ShenHai YongShi^28^. Sediment cores penetrated 18 to 20 cm into the seabed. Details for DNA sequencing can be found elsewhere^28^ and involved genomic DNA extraction with the MO BIO PowerSoil DNA Isolation Kit followed by sequencing on the MGI sequencing platforms DNBSEQ-T1 or BGISEQ500 (MGI Tech Co., Ltd., China) at BGI-Shenzhen (China).

For assessing microbial community composition, metagenomic reads were screened using singleM v0.13.2 (https://github.com/wwood/singlem) to extract *rp1B* operational taxonomic units (OTUs). Quality-control of raw reads, assembly of clean reads into contigs, and generation of metagenome-assembled genomes (MAGs) used the metaWRAP^58^ pipeline (v1.3) with details as reported previously^28^. Following depreciation using dRep v3.0.0 (parameters: -comp 50 -con 10 -sa 0.95)^59^ a non-redundant set of 487 species-level MAGs was obtained. Taxonomic classifications of bacterial and archaeal genomes were assigned using GTDB-Tk v2.1.1 with the Genome Taxonomy Database using the R207_v2 reference package^60^. The set of 120 bacterial or 53 archaeal marker genes were identified, aligned, concatenated and trimmed using GTDB-Tk v2.1.1. Genomes are then placed into the domain-specific trees using IQ-TREE v2.0.5 with best-fit models and 1000 ultrafast bootstraps^61^. Bacterial and archaeal trees were visualized and beautified in the Interactive Tree Of Life (iTOL; v6)^62^. DefenseFinder v1.0.2 (parameters: -dbtype gembase)^8^ was used to systematically detect antiviral defense systems in MAGs based on MacSyFinder models v1.2.0 in line with MacSyfinder rules^63^.

### Enumeration of viruses via fluorescence microscopy

Viral particles in sediments were counted by fluorescence microscopy according to a previous protocol^64^. In brief, around ~0.8 g sediment from each sample was transferred into a sterile 50 mL centrifuge tube and promptly fixed in 0.5% glutaraldehyde. Viruses were separated from sediments by vortexing in the dark, incubated in sodium pyrophosphate, and sonicated on ice. Samples were then filtered onto 0.02 μm pore-size membrane filters (Anodisc 25, Whatman), stained with SYBR Green I and observed using a HORIBA Aqualog fluorescence microscope (Tokyo, Japan) with a Leica imaging system. The Find maxima tool of Image J (https://imagej.net) was used to automatically select the fluorescent points^65^ with manual curation.

### Virus identification, vOTU clustering and taxonomic assignment

Potential single-contig viral genomes were identified from 18 metagenomic assemblies (contigs larger than 10 kb) using DeepVirFinder v1.0 ^66^, Virsorter2 v2.2.3 ^67^ and VIBRANT v1.2.1 ^68^. Additionally, the MetaviralSPAdes module of SPAdes v3.15.2 was used to assemble viral contigs from metagenomic reads with default parameters^69^. CheckV v1.0.1 (database v1.1)^37^ was applied to estimate the completeness and contamination of contigs identified (n=6,520) using the above four methods. Genomes with ≥50% estimated completeness (n = 488) were clustered into species-level vOTUs according to MIUViG guidelines (95% average nucleotide identity; 85% aligned fraction)^38^. Clustering used the method for single-contig viral genomes^41^ based on the supporting code of the CheckV v1.0.1 repository^15, 37^. Representative viral genomes for each species-level vOTU (n = 338) were clustered into viral clusters (VCs) that were roughly equivalent to ICTV (International Committee on Taxonomy of Viruses) prokaryotic viral genera using vConTACT2 v0.11.3 (parameters: --pcs-mode MCL --vcs-mode ClusterONE --rel-mode ‘Diamond’ --db ‘ProkaryoticViralRefSeq94-Merged’) enabled by gene-sharing networks^39^. The geNomad v1.3.3 pipeline (genomad end-to-end)^41, 70^ was employed for the taxonomic assignment of viral genomes in accordance with the taxonomy contained in ICTV’s VMR number 19 (https://ictv.global/). BACPHLIP v0.9.6 (with a minimum score of ≥ 0.8)^71^ and VIBRANT v1.2.1^68^ were used to test if complete viral genomes were likely to be either temperate (lysogenic) or virulent (lytic). Remaining viral genomes were predicted to be lysogenic or unclassified depending on if they contained provirus integration sites or integrase genes based on the annotation provided with each genome.

### Host assignments for bacteriophages and archaeoviruses

A total of 2,678 bacterial and archaeal MAGs recovered from 68 previously sequenced cold seep sediments were used to serve as the host reference database^1^. Multiple host prediction strategies were used to link viral genomes to their microbial hosts following our previous method^7^ complemented with iPHoP, an automated command-line pipeline for host predictions^72^ (**Supplementary Figure 4**). (i) For CRISPR spacer matches, the CRISPR arrays of cold seep microbial genomes were predicted using the CRISPRidentify v1.1.0 with default parameters^73^. Spacers shorter than 25 bp and CRISPR array with fewer than three spacers were dropped out. CRISPR spacers were aligned with viral genomes with ≤1 mismatch using BLASTn, and the thresholds of 95% identity were selected. Additionally, 1,398,130 spacers from 40,036 distinct genomes in the iPHoP_db_Sept21 database were also used for CRISPR-based predictions by version 1.1.0 of iPHoP^72^. (ii) For the detection of shared tRNA between viruses and hosts, tRNA genes were annotated using tRNAscan-SE v2.0.9 (parameters: -B -A)^74^. Putative host-virus linkages satisfied a threshold of ≥90% length identity over the 95% of the sequences by BLASTn. (iii) For alignment-based matches, viral genomes were aligned with microbial genomes using BLASTn based on their nucleotide sequence homology (e-value ≥ 0.001, nucleotide identity ≥ 70%, match coverage over the length of viral genomes ≥ 75% and bitscore ≥ 50). (iv) For host predictions based on independent signals (k-mer usage profiles and protein content), VirHostMatcher (VHM)^75^, WIsH^76^, Prokaryotic virus Host Predictor (PHP)^77^ and RaFAH^78^ were performed individually using iPHoP v1.1.0. Match criteria were d_2_* values ≤ 0.2 for VHM, *p* values ≤ 0.05 for WIsH, the predicted ‘maxScoreHost’ for PHP and RaFAH_scores > 0.14 for RaFAH. The genome was considered to be the host if it belonged to the same family with top hits for each viral genome based on multiple methods. When multiple hosts from different phyla for a viral genome were predicted according to the four methods, all these virus-host linkages were ignored.

### Identification of counter-defense systems, reverse transcriptases, auxiliary metabolic genes and alternative genetic codes

For counter-defense systems, Acr-Aca operons were predicted based on the guilt-by-association approach using Acafinder (--Virus; version of Oct 15, 2022)^42^. Restriction-modification (R-M) systems were identified using previous Hidden Markov model profiles and scripts (https://github.com/oliveira-lab/RMS; version of Mar 16, 2018)^34^. Toxin and Antitoxin (TA) genes were identified based on specific domains of TA systems using Metafisher (https://github.com/JeanMainguy/MeTAfisher). Reverse transcriptases (RTs) were predicted and classified through the myRT web-server (https://omics.informatics.indiana.edu/myRT/)^14^.

Auxiliary metabolic gene (AMG) identification was performed following previous protocols^7, 79^. Briefly, checkV-trimmed viral genomes were run through VirSorter2 (--prep-for-dramv) to produce the viral-affi-contigs-for-dramv.tab, and then the annotations were done using DRAM v1.2.0 (viral mode; default parameters)^80^. Genes with auxiliary scores of 1-3 and AMG flags of M and F were considered putative AMGs for further validation by manual checking of gene locations. PROSITE^81^ was used to analyze the conserved domains of AMGs and SWISS-MODEL^82^ was used for protein structural predictions. Three-dimensional structures of viral AMGs were predicted using ColabFold by combining the fast homology search of MMseqs2 with AlphaFold2 ^83, 84^. Genome maps of AMG-containing viral genomes were visualized based on DRAM-v annotations using Easyfig v.2.2.0 (ref.^85^).

Mgcod v1.0.0 was used to identify blocks with specific genetic codes for cold seep viral genomes (parameters: --isoforms)^86^. In this pipeline, MetaGeneMark^87^ was applied to find the highest scoring model among four genetic code models: i) the standard genetic code (genetic code 11), ii) a model with the *opal* (TGA) reassignment (genetic code 4), iii) a model with the *amber* (TAG) reassignment (genetic code 15), and iv) a model with the *ochre* (TAA) reassignment (genetic code 101). Identified recoded regions were annotated using eggnog-mapper v2.1.9 (ref.^88^) against the eggNOG database (v5.0)^89^.

### Macro- and microdiversity analyses of viral populations

Filtered reads from each sample were mapped to 338 single-contig viral genomes that represent each vOTU using Bowtie2 v 2.3.5 ^90^. Resulting BAM files, viral genomes, and read counts for each metagenome were used as input for the MetaPop pipeline^91^ for pre-processing, macrodiversity and microdiversity analyses. MetaPop was run using the default parameters (--snp_scale both) and genes from viral genomes were predicted using Prodigal v2.6.3 ^92^. Macrodiversity estimates include population abundances, alpha-diversity (within community) and beta-diversity (between community) indices. To accurately call SNPs and assess contig-level microdiversity, 207 viral populations with >10× average read depth coverage and >70% length of genome covered were retained for microdiversity analyses^91^. SNP frequencies subsampled down to 10× coverage were used to assess nucleotide diversity *(θ* and *π*) at the individual gene and whole-genome levels, as well as fixation indices (F_ST_; between population microdiversity) and selective pressures on specific genes (pN/pS and Tajima’s D).

### Statistical analyses

Statistical analyses were performed using R v4.0.0. The normality and variance homogeneity of the data were assessed using Shapiro-Wilk and Bartlett’s tests. Wilcoxon tests were used to compare differences in viral microdiversity parameters (π, Tajima’s D, pN/pS) across cold seep stages. The Kruskal-Wallis rank-sum test with Chi-square correction was used to compare differences in evolutionary metrics of genomes and genes among different groups and samples. Correlations between microdiversity and sediment depth, defense system numbers, genome sizes, and others parameters were obtained using the linear regression with the fitness and confidence of the regression curves characterized as R-square and p values, respectively.

## Data availability

MAGs, vOTUs, AMGs and other related information have been uploaded to figshare (https://figshare.com/s/xxx).

## Acknowledgements

The work was supported by the Scientific Research Foundation of Third Institute of Oceanography, MNR (No. 2022025), the National Key Research and Development Program of China (2022YFC2805500), State Key Laboratory of Marine Geology, Tongji University (No. MGK202303), and China Postdoctoral Science Foundation (2022M723709). We thank Dr. Xiaoyuan Feng for providing valuable comments.

## Author contributions

X.D. designed this study. X.D., Y.P., and Z.L. performed the omics analysis. Z.Z. and Z.L. counted virus particles in sediments. X.D., Y.P., P.D., L.-D.S., Q.L., C.Z., K.J., and C.R.J.H. interpreted the data. J.L. contributed to data collection. X.D., Y.P., and C.R.J.H. wrote the paper, with input from other authors.

## Conflict of interest

The authors declare no competing interest.

## Notes

### Competing Interest Statement

The authors have declared no competing interest.

### Summary of Updates

author affiliations updated

