## Supplementary Figures for "Viruses in deep-sea cold seep sediments harbor diverse survival mechanisms and remain genetically conserved within species"

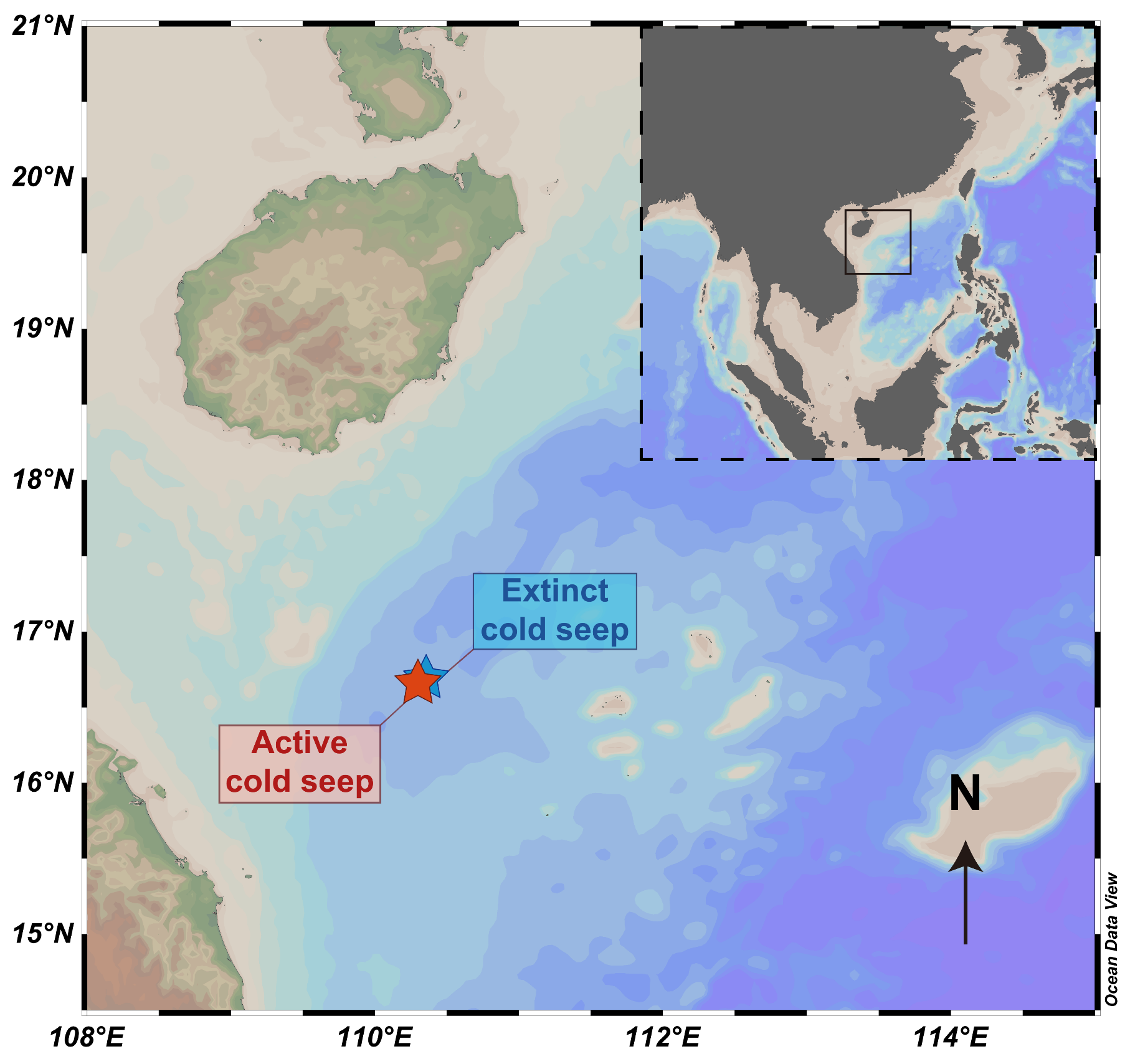
**Supplementary Figure 1. Location of the Haima cold seep study area**. Sampling locations of the active and extinct cold seep sites in the Haima cold seep are indicated. This map is drawn using the Ocean Data View v5.4.0.

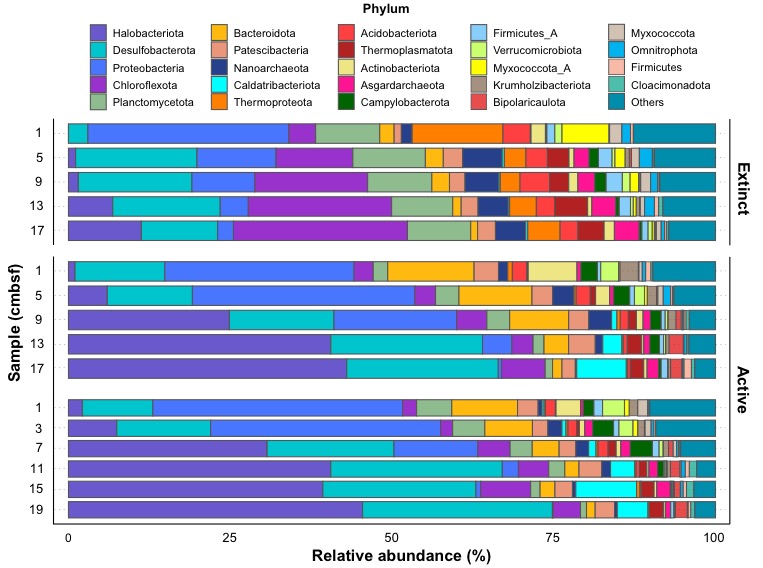
**Supplementary Figure 2. Phylum-level relative abundance of taxa in different cold seep sediment types (active and extinct).** Comparison of taxonomic differences in microbial communities of the 16 samples along the sampling sites and vertical depths based on *rplB*. The diagram shows the top 25 microbial phyla with the highest relative abundance. Detailed data for the relative abundance of each population in each sample can be found in **Supplementary Table 2**.

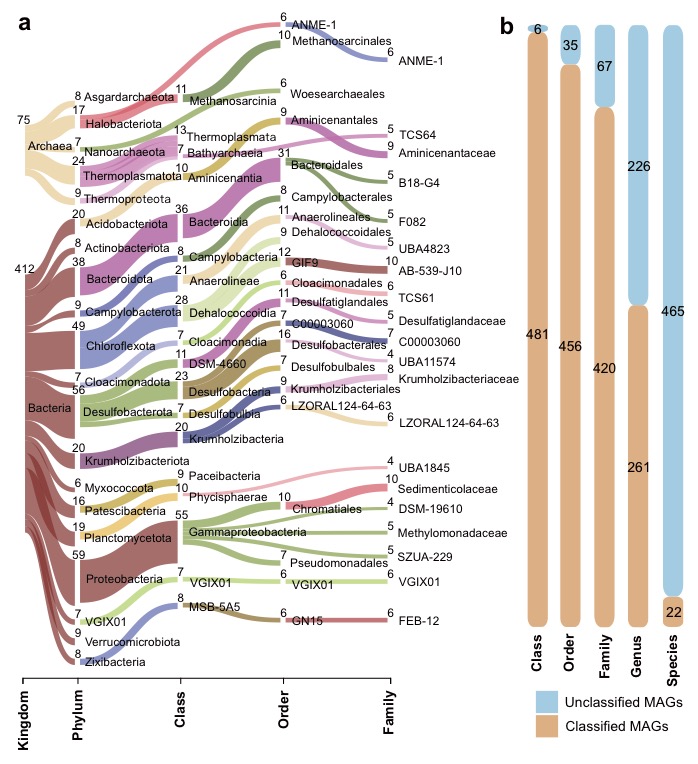
**Supplementary Figure 3. Classification of MAGs recovered from Haima active and extinct cold seep sediments.** (a) Sankey diagram based on assigned GTDB taxonomy showing recovered archaeal and bacterial MAGs at different taxonomic levels, indicating the number of MAGs recovered for a given lineage. The diagram shows the top 25 taxa with the largest number of MAGs at each level. (b) Proportion of MAGs classified by GTDB-Tk at each taxonomic level. Detailed data for the information of each MAG can be found in **Supplementary Table 3**.

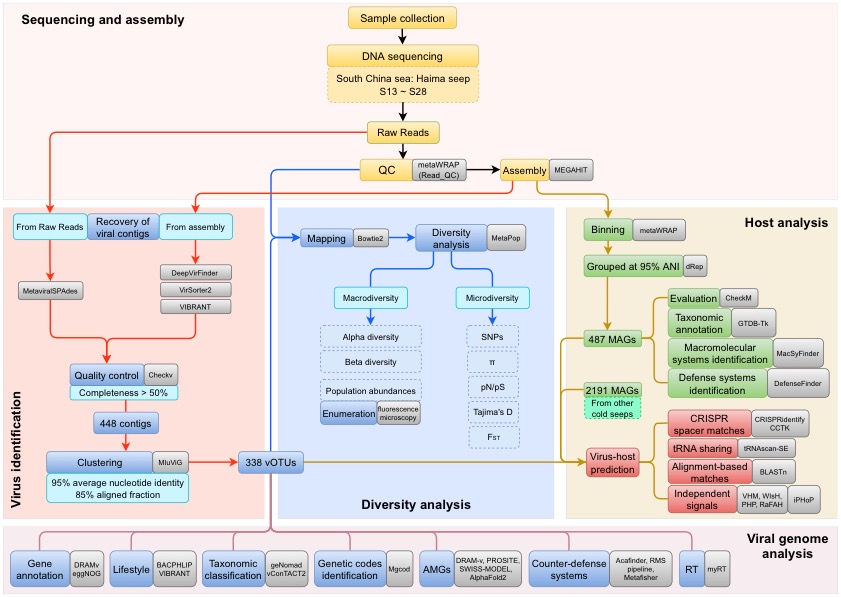
**Supplementary Figure 4. Bioinformatics workflow for analysis of viral populations.** The process consists of five modules: sequencing and assembly, virus identification, diversity analysis, host analysis, and viral genome analysis. Yellow: data collection; green: microbial analysis; blue: viral analysis; red: interactions between hosts and viruses; grey: software or equipment used.

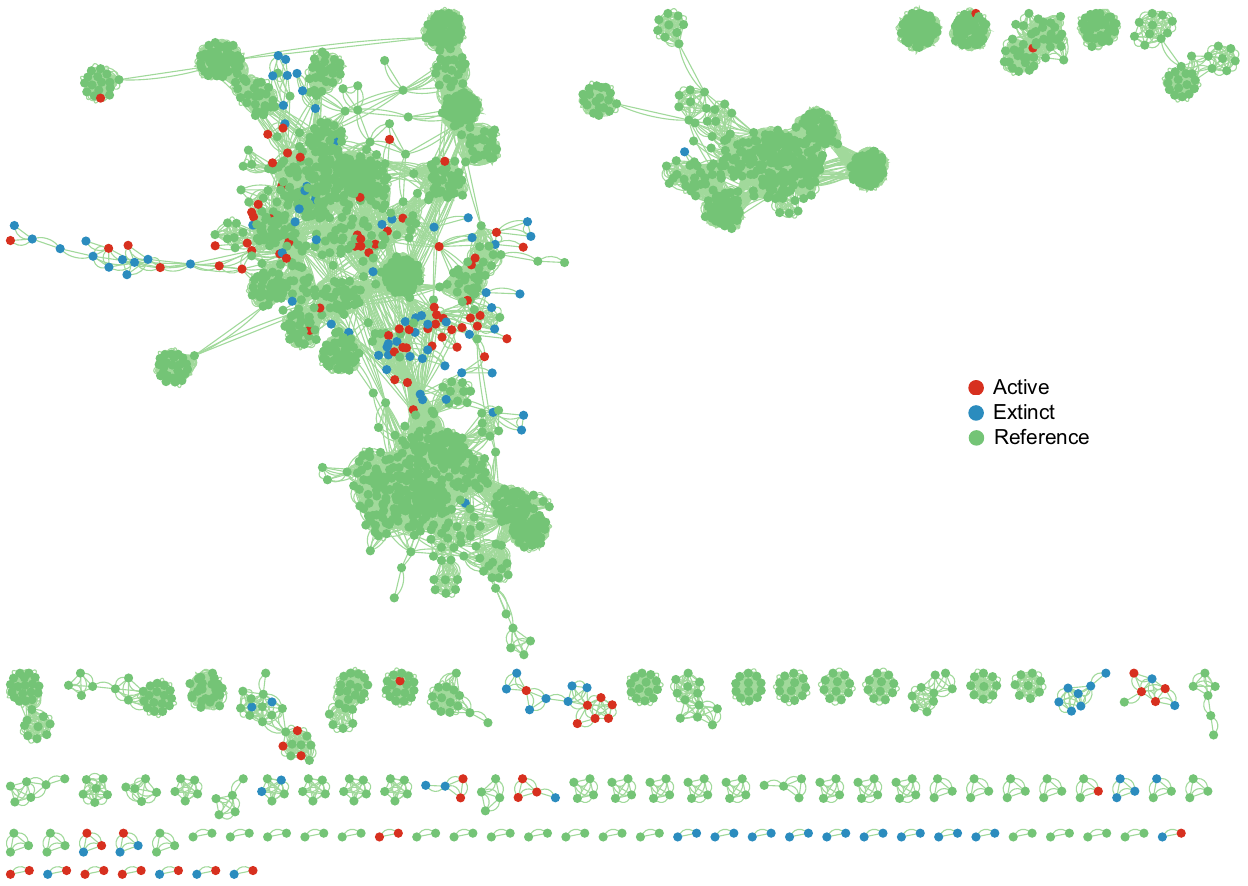
**Supplementary Figure 5. Viral contig clustering in cold seep sediments.** Gene-sharing network of viral sequence space based on assembled viral genomes from cold seep sediment and RefSeq prokaryotic viral genomes (v94). Nodes represent viral genomes and edges indicate similarity based on shared protein clusters. Detailed information for clustered vOTUs is provided in **Supplementary Table 6**. The gene-sharing network is visualized in Cytoscape v3.9.1.

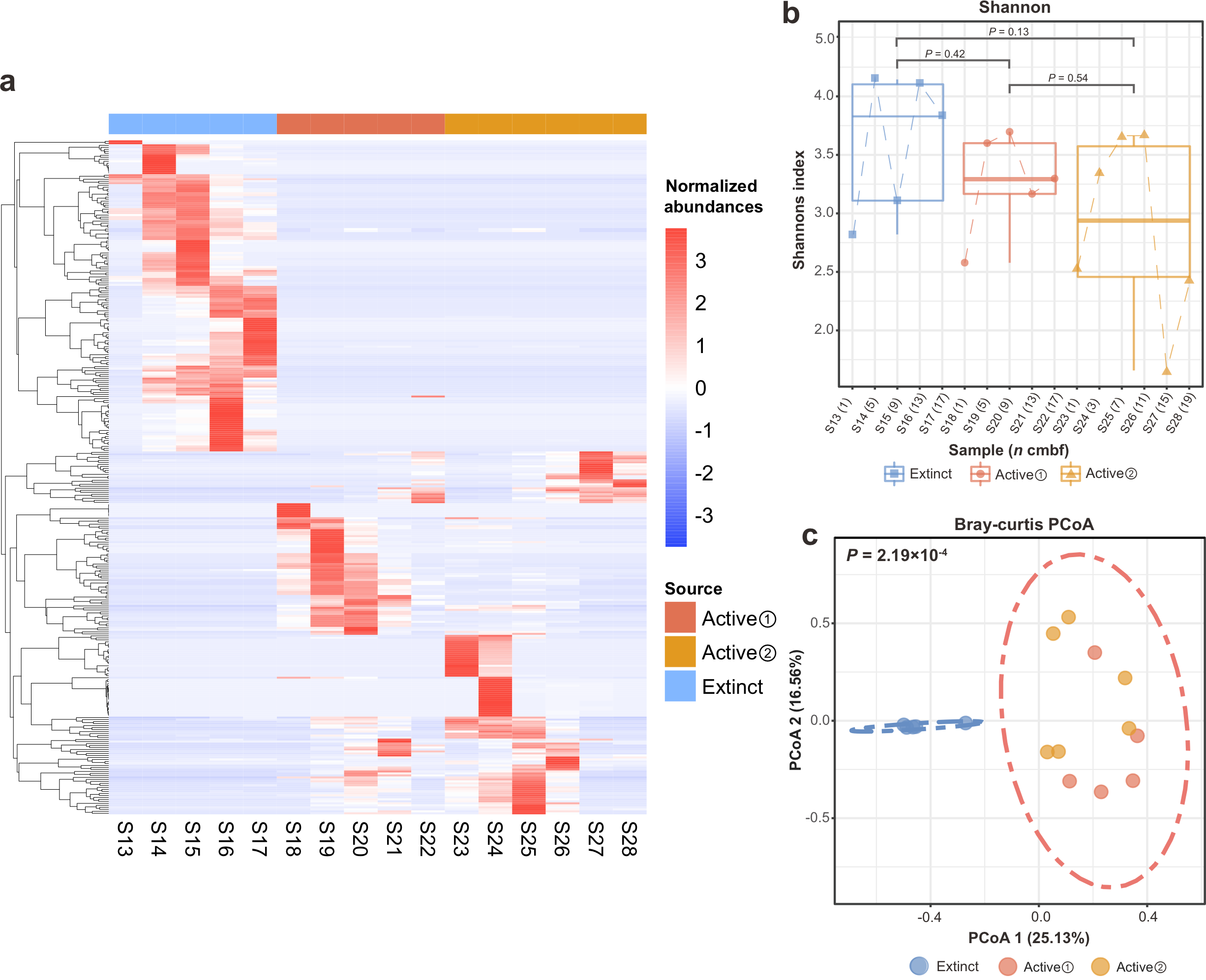
**Supplementary Figure 6. Normalized abundances of viruses and viral diversity in sediments from three cold seep sites.** (a) Columns indicate the 16 different sediment samples from the three cold seep sites. The dendrogram (at left) and rows represent the 338 vOTUs recovered from cold seep sediment metagenomes. The color intensity of each cell in the resulting matrix reflects the normalized abundance of a given genome in a given sample. (b) Shannon index of vOTUs for each sediment sample. The values in parentheses for the column labels represent the sediment depth (cmbsf) for each sample. P-values of differences across different cold seep sites were calculated using a Wilcoxon test. (c) Bray-Curtis dissimilarity calculated from normalized abundances of vOTUs. ADONIS was applied to test the difference in viral communities between samples from active and extinct cold seep phases. The p-value indicates highly significant differences in viral diversity between active and extinct cold seep sediments, which can be separated by 95% confidence ellipses. Detailed data for the normalized abundance and Shannon index of vOTUs can be found in **Supplementary Table 5**.

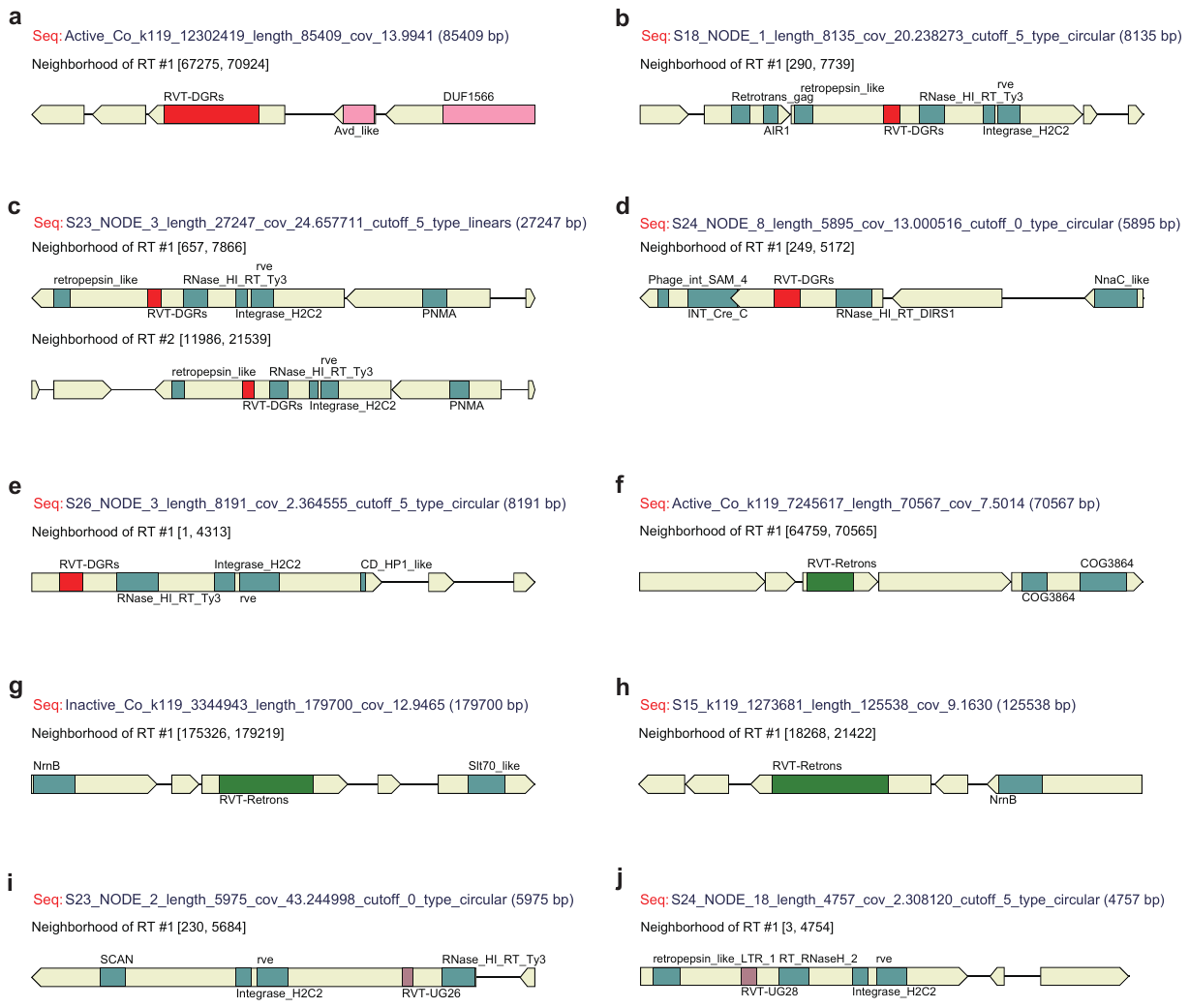
**Supplementary Figure 7. Reverse transcriptases (RTs) in cold seep viruses are predicted using MyRT.** Loci of predicted RT genes and its neighborhood along the genome are shown. Arrows represent the genes, with the different regions encoding different domains in colored rectangles. RTs of different classes are in different colors (RVT-DGRs, red; RVT-Retrons, green; RVT-UG26 and RVT-UG28, brown).

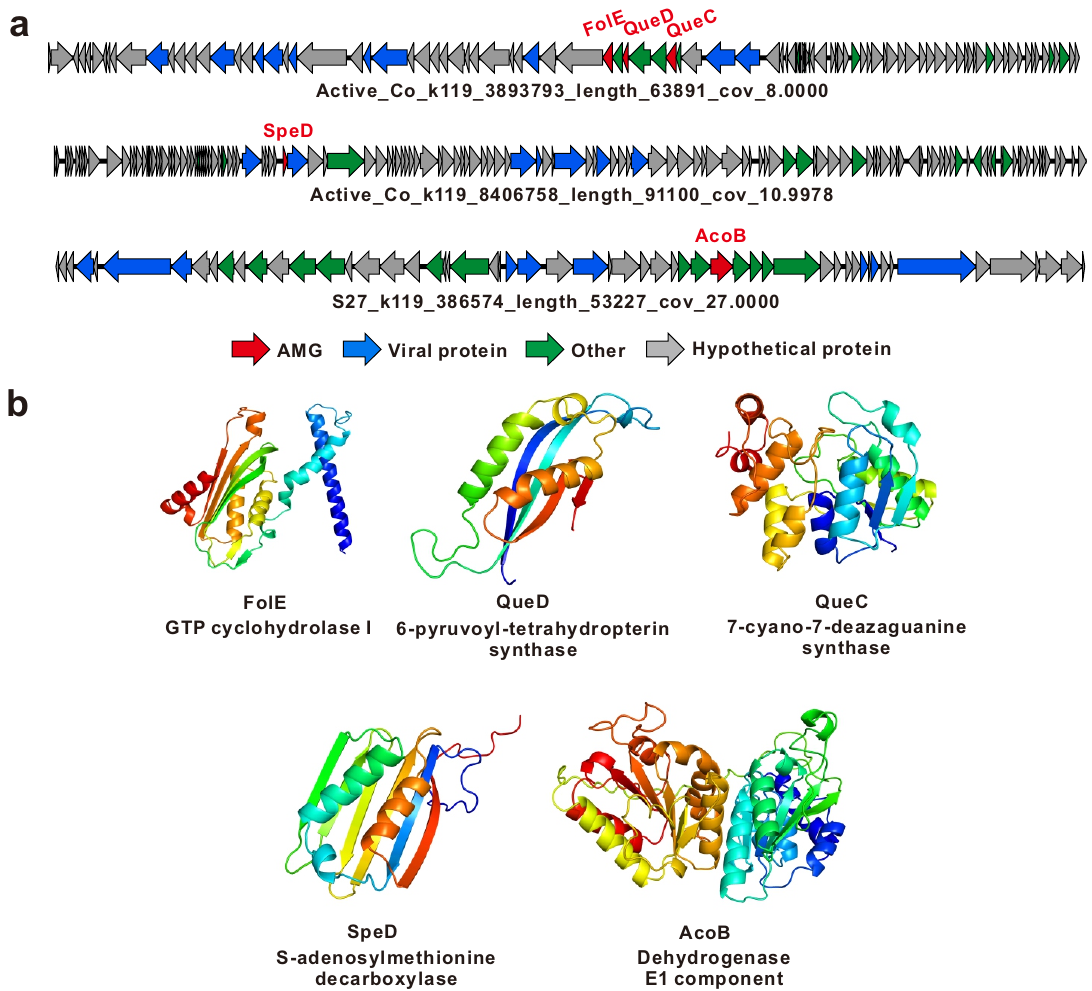
**Supplementary Figure 8.** **Genome synteny across viruses with auxiliary metabolic genes (AMGs) and their structural predictions.** (a) Genome map of three viral genomes encoding five AMGs (indicated in red) together with conserved core viral genes (indicated in blue). (b) AMG structural predictions for FolE, QueD, QueC, SpeD and AcoB using Alphafold2 online. Detailed statistics for virus-encoded AMGs are provided in **Supplementary Table 9**.

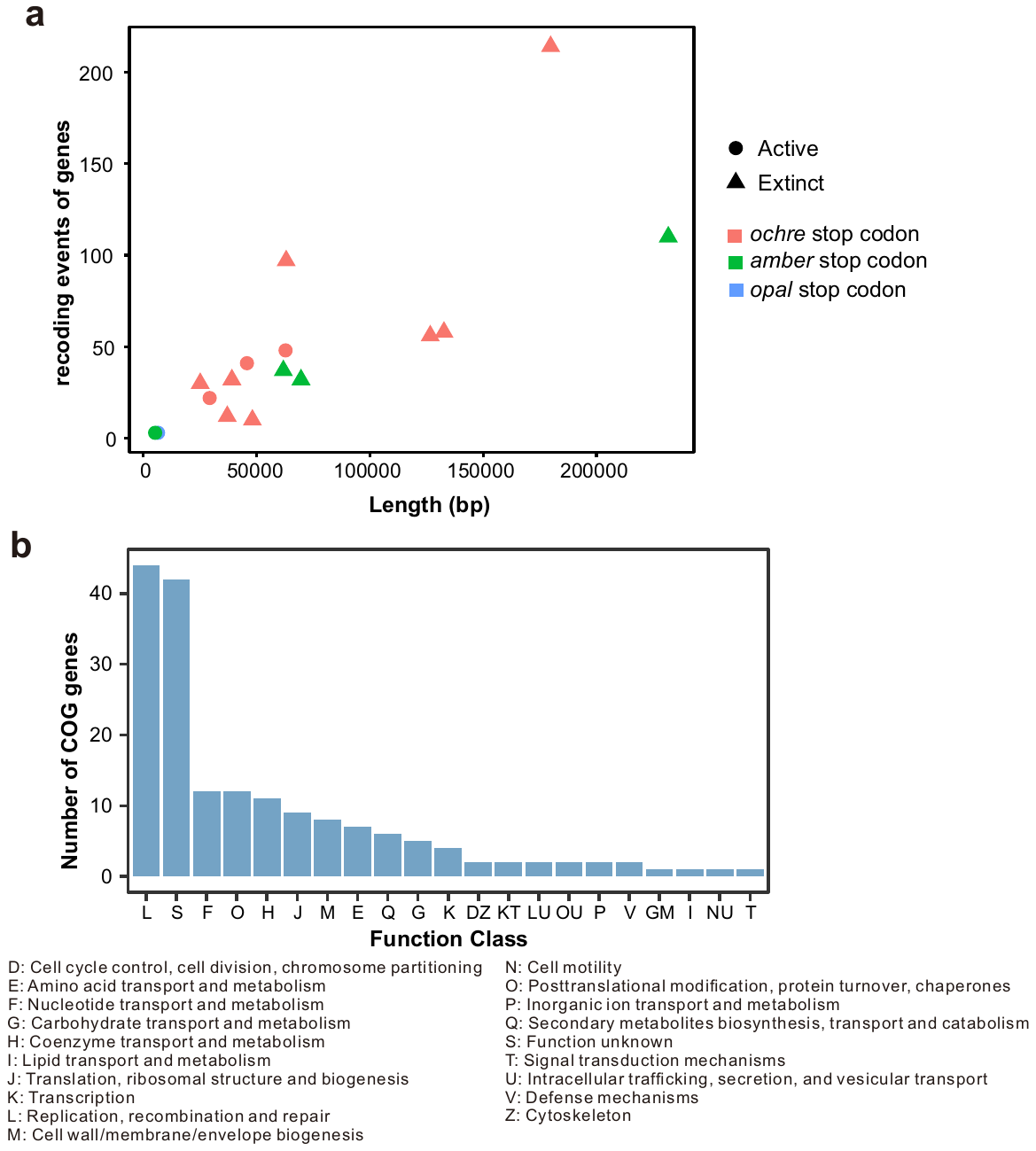
**Supplementary Figure 9. Alternative genetic code assignments observed in cold seep viruses.** (a) Correlation between numbers of recoding events of genes with multiple genetic code assignments and genome size. (b) Classification of viral genes with multiple genetic codes into COG functional categories. Detailed statistics for alternative genetic coding of viruses are provided in **Supplementary Table 10**.

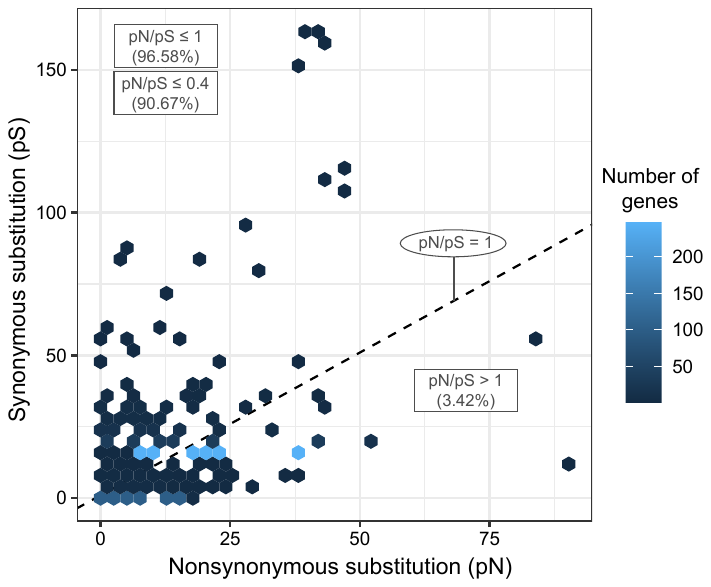
**Supplementary Figure 10. Comparison of synonymous and nonsynonymous substitutions of viral genes in cold seep sediments.** The position of the dashed line indicates pN/pS = 1. Detailed statistics for microdiversity of viral genes are provided in **Supplementary Table 13**.

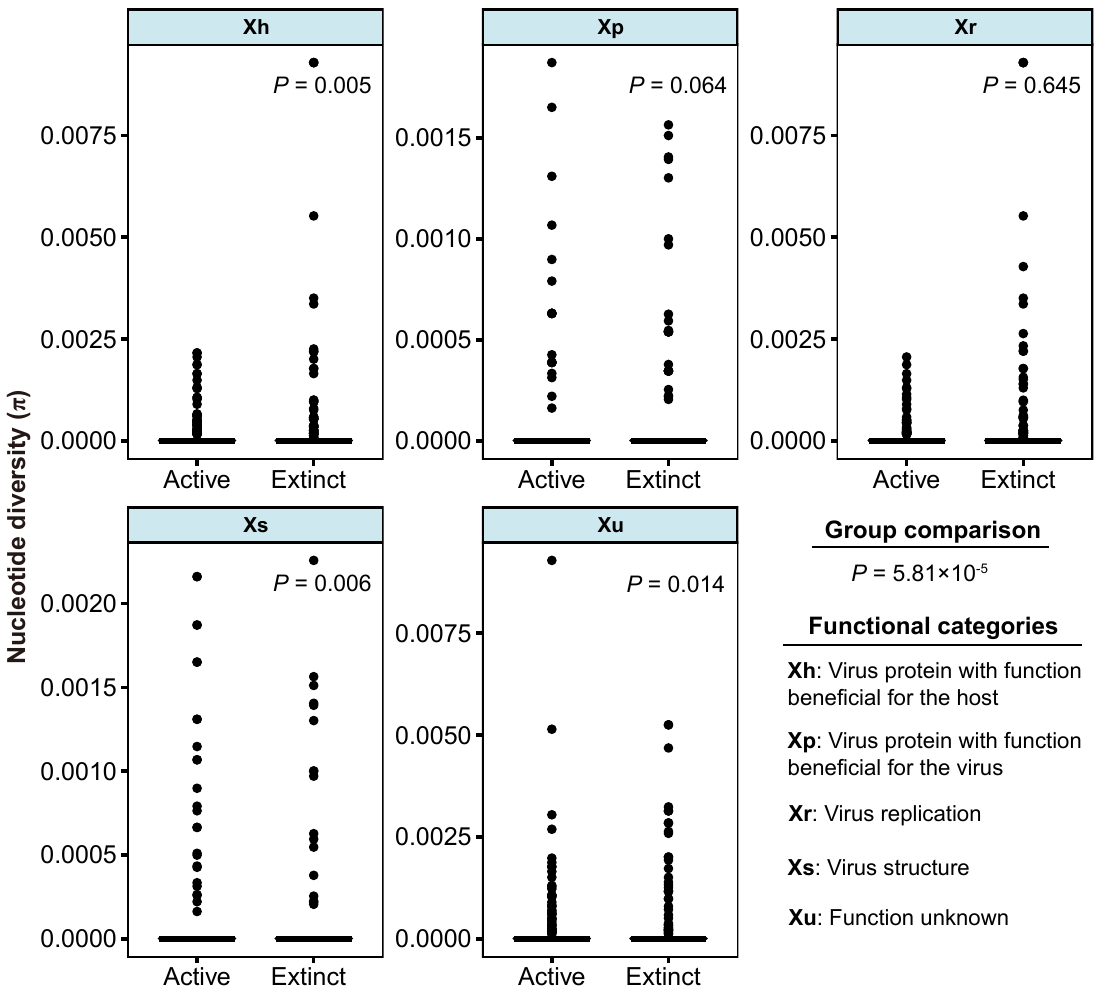
**Supplementary Figure 11. Microdiversity of viral genes grouped according to the functional categories.** Comparison of nucleotide diversity of viral genes (virus replication, virus structure, protein with function beneficial for the host or the virus and function unknown) in active and extinct cold seep sediments. Detailed data on microdiversity of viral genes can be found in **Supplementary Table 13**.
